# hERG1 channel subunit composition mediates proton inhibition of IKr in hiPSC-CMs

**DOI:** 10.1101/2022.08.26.505466

**Authors:** Chiamaka U. Ukachukwu, Eric N. Jimenez-Vazquez, Abhilasha Jain, David K. Jones

## Abstract

hERG1 conducts cardiac IKr and is critical for repolarization of the human heart. Reduced IKr causes long QT syndrome and increases the risk for cardiac arrhythmia and sudden cardiac death. At least two subunits combine to form functional hERG1 channels, hERG1a and hERG1b. Changes in hERG 1a/1b subunit abundance modulates IKr kinetics, magnitude, and drug sensitivity. Studies from native cardiac tissue have suggested that hERG1 subunit abundance is dynamically regulated, but the impact of altered subunit abundance on IKr and its response to external stressors is not well understood. Here, we used a substrate-driven hiPSC-CM maturation model to investigate how changes in relative hERG 1a/1b subunit abundance impact the response of native IKr to extracellular acidosis, a known component of ischemic heart disease and sudden infant death syndrome. IKr recorded from immature hiPSC-CMs display a two-fold greater inhibition by extracellular acidosis (pH 6.3) compared to matured hiPSC-CMs. qRT-PCR and immunocytochemistry demonstrated that hERG1a subunit mRNA and protein were upregulated, and hERG1b subunit mRNA and protein were downregulated in matured hiPSC-CMs compared to immature hiPSC-CMs. The shift in subunit abundance in matured hiPSC-CMs was accompanied by an increased in IKr density. Silencing the impact of hERG1b on native IKr kinetics by overexpressing a polypeptide identical to the hERG1a PAS domain reduced the magnitude of IKr proton inhibition in immature hiPSC-CMs to levels comparable to those observed in matured hiPSC-CMs. These data demonstrate that hERG1 subunit abundance is dynamically regulated and that hERG1 subunit abundance determines IKr sensitivity to protons in hiPSC-CMs.

## INTRODUCTION

hERG1, encoded by KCNH2, is the voltage-gated potassium channel that conducts the rapid delayed rectifier potassium current (IKr). Reduced IKr from either off-target pharmacological block or loss-of-function KCNH2 variants causes the cardiac disorder long QT syndrome and increases the risk for cardiac arrhythmia, syncope, and sudden cardiac death (1,2). Long QT syndrome is the leading cause of arrhythmic death in children and accounts for 5-10% of sudden infant death syndrome (SIDS) and intrauterine fetal death cases (3-8). Furthermore, multiple LQTS-associated KCNH2 variants have been linked with intrauterine fetal death and SIDS, underscoring the importance of hERG1 in the young heart (3,9-13).

At least two hERG1 subunits comprise native hERG1 channels, hERG1a and hERG1b (14-18). Mutations in both subunits promote/ cause cardiac electrical dysfunction (3,19-21). hERG1a subunits contain an N-terminal Per-Arnt-Sim (PAS) domain that regulates channel gating through interactions with the C-terminal cyclic nucleotide binding homology domain (CNBHD) and the cytoplasmic S4-S5 linker (20,22-25). hERG1b has a much shorter and unique N-terminus that lacks a PAS domain (14,15). When heterologously expressed in HEK293 cells, the absence of a functional PAS domain in hERG1b triggers a roughly two-fold acceleration in the time course of activation, deactivation, and inactivation recovery in heteromeric hERG 1a/1b channels compared to homomeric hERG1a channels (20). In cardiomyocytes derived from human induced pluripotent stem cells (hiPSC-CMs), silencing hERG1b by overexpressing a polypeptide that mimics the hERG1a PAS domain slows native IKr gating kinetics and reduces IKr magnitude, triggering increased action potential duration and early afterdepolarizations (16). Conversely, disabling the hERG1a PAS domain using PAS-targeting antibodies accelerates IKr gating, increases IKr magnitude, and hastens cardiac repolarization (26).

Extracellular acidosis is a major inhibitor of IKr (27), and occurs in a variety of pathological situations associated with cardiac dysfunction, including SIDS and myocardial ischemia (28-30). Consequently, a large body of work has explored the impact of extracellular protons on hERG1 (27,31-37). Briefly, reduced extracellular pH reduces hERG1 channel conductance, depolarizes channel voltage dependence, and accelerates channel deactivation (31,33,34,38). The pro-arrhythmic effects of reduced pH on hERG1 are two-fold, pore block by protons slows cardiac repolarization whereas the accelerated deactivation impairs hERG1’s ability to protect the heart from premature stimulation (27,39,40). Interestingly, it was demonstrated that the inhibitory effect of extracellular protons is enhanced in hERG1b-containing channels (41,42).

Several studies suggest that hERG1 subunit abundance is dynamically regulated *in vivo* (18,43-48). However, the mechanisms that determine hERG1 subunit abundance and the impact of altered subunit abundance on the susceptibility of arrhythmia are poorly understood. LQTS mutations in the hERG1a PAS domain were shown to disrupt hERG1b trafficking to the membrane (49). In murine tissue, targeted mERG1b deletion abolishes IKr in adult mice but only reduces IKr magnitude by roughly 50% in neonates, compared to wildtype littermate controls (50). These data suggest that mERG1a is selectively downregulated during maturation of the murine heart. In the human heart, hERG1a mRNA transcripts are upregulated and hERG1b transcripts downregulated in adult ventricular tissue compared to fetal cardiac tissue (3). Similarly, the relative abundance of hERG1a to hERG1b protein was reduced in failing cardiac tissue compared to non-diseased donor controls (51).

In this study, we used *in vitro* maturation of hiPSC-CMs to probe the impact of hERG1 subunit dynamics on proton modulation of native cardiac IKr. The data presented herein demonstrate that increased hERG1a and reduced hERG1b in matured hiPSC-CMs diminish IKr sensitivity to extracellular protons compared to IKr recorded from immature hiPSC-CMs.

## RESULTS

Protons decrease hERG1 current amplitude and accelerate the time course of hERG1 deactivation (27,31,33,34,37,54). However, the specific effects of extracellular acidosis can vary across expression systems. For example, the impact of protons on the voltage dependence of activation is not consistently reported. These variations across systems suggest that other unidentified factors contribute to the response of hERG1 to protons (33,35-37,39,54,55). Subunit abundance is one factor that may explain the different acidosis sensitivities. To determine the impact of hERG1 subunit abundance on native IKr sensitivity, we cultured hiPSC-CMs on two different matrices to promote distinct stages of maturation and corresponding shifts in hERG1 subunit expression.

### Extracellular matrix mediates hiPSC-CM maturation

Culturing hiPSC-CMs on a pliable substrate promotes hiPSC-CM maturation (56-58). We cultured hiPSC-CMs on either a pliable substrate (polydimethylsiloxane, PDMS) or a stiff substrate (glass). All substrates were coated with Matrigel® prior to hiPSC-CM plating. Previous reports using PDMS as a substrate, hiPSC-CMs have more mature electrophysiological features (e.g., increased INa and IK1, faster upstroke velocity and faster conduction velocity, hyperpolarized RMP, etc.) compared to hiPSC-CMs plated on a hard substrate (56). Here, we found that hiPSC-CMs cultured on Matrigel-coated PDMS displayed electrophysiological characteristics consistent with enhanced maturation compared to hiPSC-CMs cultured on Matrigel-coated glass coverlips (Fig. 1). Action potentials recorded from hiPSC-CMs cultured on PDMS displayed hyperpolarized resting membrane potentials and larger action potential amplitudes compared to action potentials recorded from hiPSC-CMs cultured on glass (Fig. 1A-C). Additionally, E-4031-sensitive currents, which are indicative of native IKr, showed a trend to be increased in PDMS-cultured hiPSC-CMs compared to glass-cultured hiPSC-CMs. Steady-state IKr density, measured at the end of a three second step pulse, was increased from 1.3 ± 0.1 pA/pF in glass-cultured hiPSC-CMs to 1.9 ± 0.3 pA/pF in PDMS-cultured hiPSC-CMs (Fig. 1D-F). Tail IKr was similarly increased, from 1.4 ± 0.1 pA/pF in glass-cultured hiPSC-CMs to 2.3 ± 0.3 pA/pF in PDMS-culture hiPSC-CMs (Fig. 1D,G,H). In contrast, hiPSC-CM maturation had no effect on the voltage dependence of IKr activation (Suppl. Fig. S1). Importantly, there was no significant difference in cell capacitance between immature and matured cells (Suppl. Fig. S2), indicating that cell size was not a factor in the recorded current density.

**Figure. 1.**
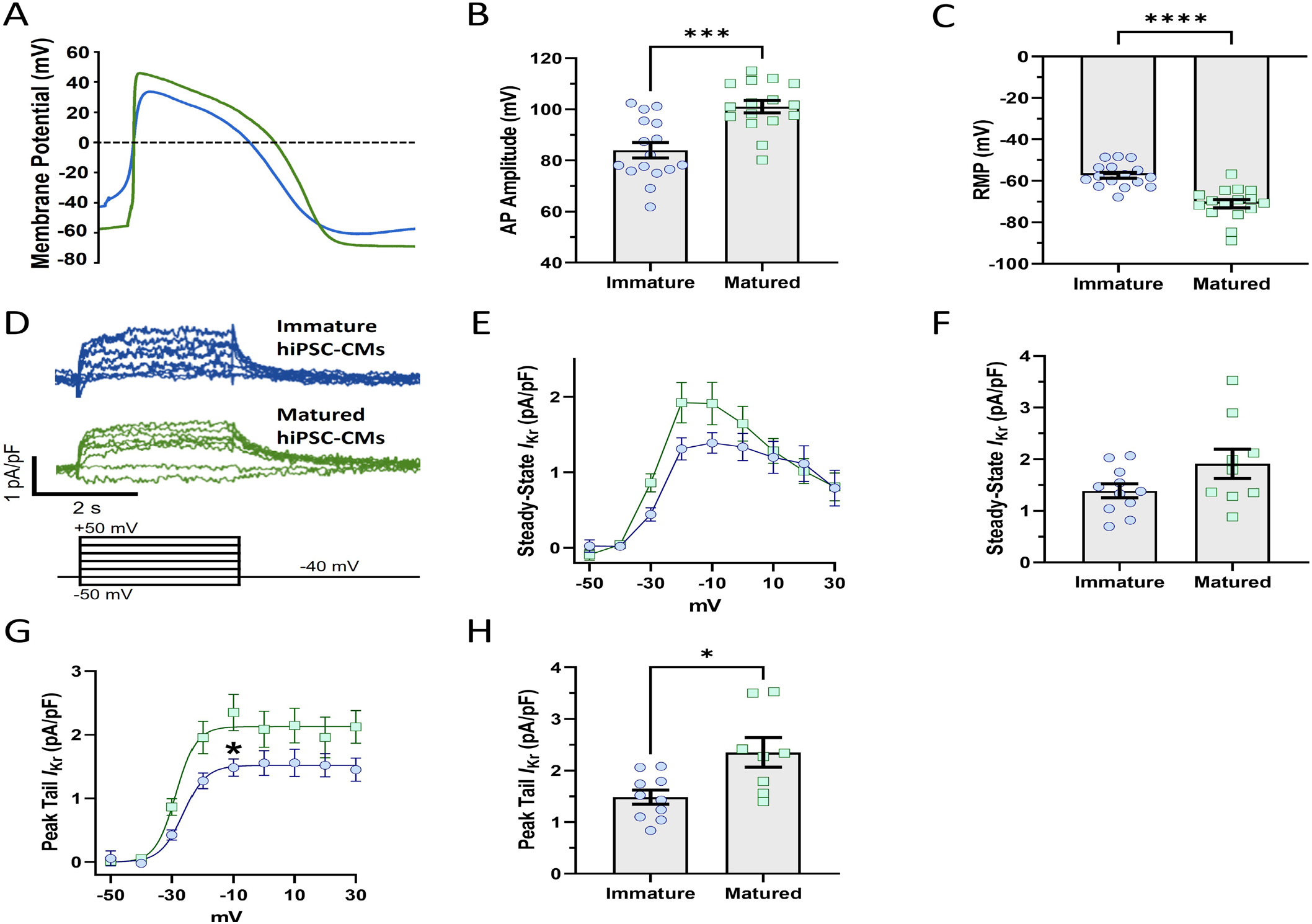
hiPSC-CM maturation with PDMS hyperpolarizes the AP and increases IKr density. **A)** Action potential recordings from cells cultured on glass (blue) and PDMS (green). **B–C)** AP parameters. Cells plated on PDMS demonstrated more hyperpolarized resting membrane potential and greater action potential amplitude than cells plated on glass. **D)** Representative IKr traces elicited by the protocol at bottom. **E)** Steady-State IKr measured at the end of the step pulse, recorded from immature and matured hiPSC-CMs. **F)** Steady-State current densities at 10 mV. **G)** Tail IKr in immature and matured hiPSC CMs. **H)** Tail current densities at 10 mV. hiPSC-CMs plated on PDMS had larger ERG currents than hiPSC-CMs plated on glass. Data were compared using a two way ANOVA test and a two tailed Mann-Whitney test where appropriate. Errors bars represent SEM. N-value = 3, n value ≥ 8. ****P < 0.0001, ***P = 0.0002, and *P < 0.05.

We also investigated the impact of hiPSC-CM maturation on IKr kinetics. We fit the decay of tail currents at -40 mV with a bi-exponential equation (Equation 2). The fits yielded fast (τfast) and slow (τslow) time constants that were similar in matured (118.4 ± 10 ms and 1,173 ± 216 ms for τfast and τslow, respectively) compared to immature hiPSC-CMs (110 ± 15 ms and 1313.5 ± 174 ms for τfast and τslow, respectively) (Suppl. Fig. S3A,B). We also recorded IKr during a voltage command designed to mimic a human ventricular action potential (Suppl. Fig. S3C). We integrated E 4031-sensitive currents elicited during the AP waveform and normalized the resultant charge to cell capacitance. Surprisingly, despite the substantial increase in tail IKr density, there was no significant difference in IKr charge densities recorded from immature and matured hiPSC-CMs (Suppl. Fig. S3D). To test if additional changes in IKr kinetics could be present in matured hiPSC-CMs, we normalized the IKr charge recorded during the AP waveform to the peak tail IKr recorded from the same cell. Like IKr deactivation, relative repolarizing charge in matured hiPSC-CMs trended to a reduction compared to immature hiPSC-CMs, but the difference was not statistically significant (P = 0.22, Suppl. Fig. S3E). This may indicate differences in gating kinetics, where channel activation is slowed, or inactivation stabilized.

### External acidosis differentially impacts IKr recorded from mature and immature hiPSC CMs

Acidosis has complex electrophysiological effects on hERG1 channels that lead to altered electrical activity. The effects of acidosis on IKr have been studied previously, revealing changes in the voltage-dependence of activation when the pH was adjusted from 7.4 to 6.3 (27). Here, we studied the impact of extracellular acidosis on native IKr recorded from either immature or matured hiPSC-CMs (Fig. 2). Figure 2A depicts representative paired IKr traces, recorded first in bath solution titrated to pH 7.4 then bath solution titrated to pH 6.3. For IKr recorded from either immature or matured hiPSC CMs, pH 6.3 decreased the step pulse and tail pulse current density by ∼50% (Fig. 2B-E), depolarized the voltage-dependence of activation by ∼12 mV (Fig. 2F), and dramatically accelerated the time course of deactivation (Fig. 2G).

**Fig. 2.**
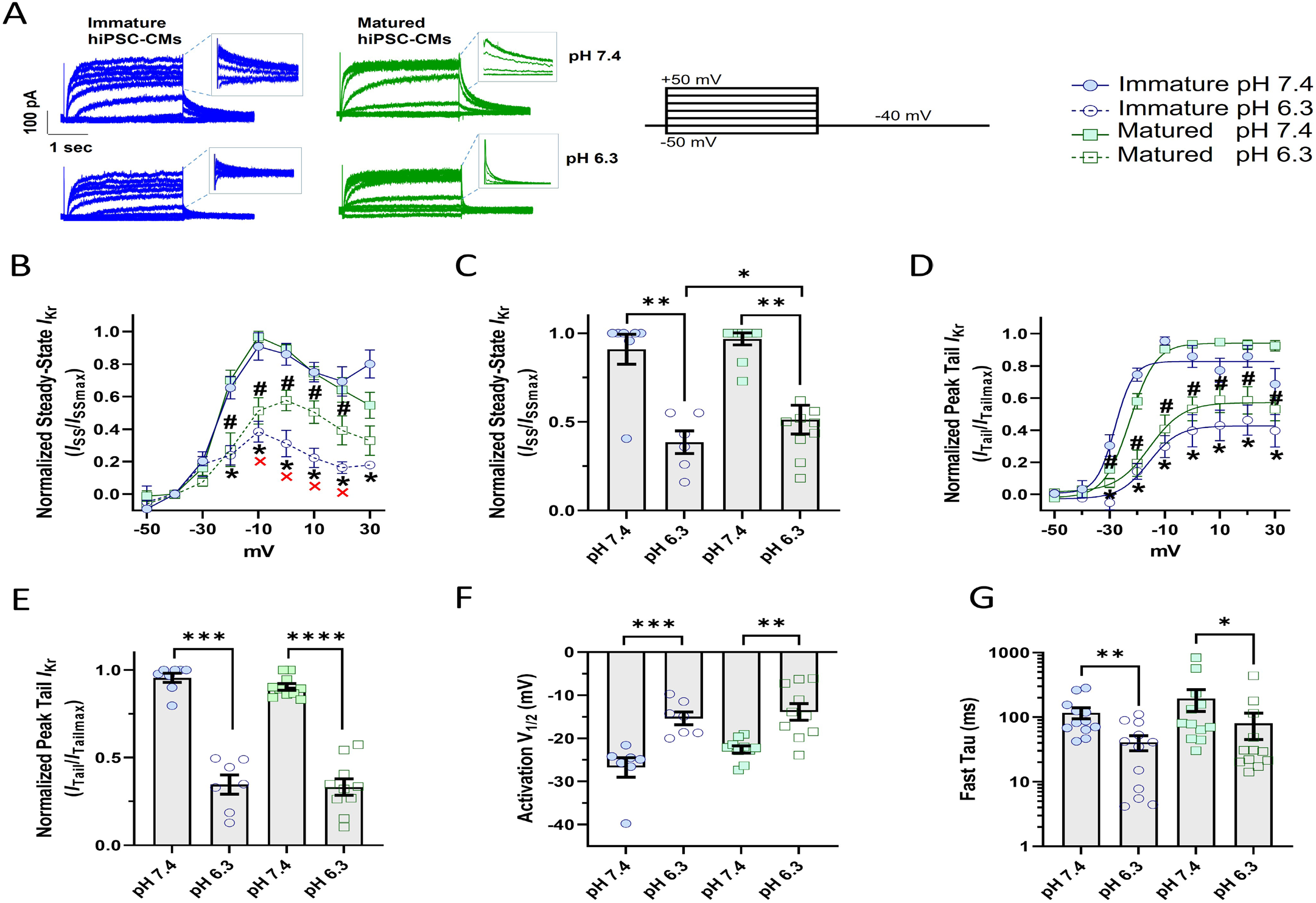
Proton sensitivity of native IKr corresponds with hiPSC-CM maturation. **A)** Representative IKr traces elicited by the protocol below from matured and immature hiPSC-CMs at pH 7.4 and pH 6.3. **B)** Steady-state IKr density in immature and matured hiPSC-CMs at pH 7.4 and pH 6.3. **C)** Normalized Steady-State current densities at 10 mV. **D)** Steady-state and peak-tail IKr density in immature and matured hiPSC-CMs in control and acidic environment. **E)** Normalized peak-tail IKr densities at 10 mV. The symbols * and # represent the statistical significance of normalized Peak tail and Steady-State IKr at pH 7.4 vs pH 6.3 in immature and matured hiPSC-CMs, respectively. The symbol × denotes significant difference between immature and matured hiPSC-CMs at pH 6.3. **F)** Voltage-dependence of activation (V1/2) for IKr from immature and matured hiPSC-CMs in control and acidic environment. **G)** Time constants of IKr deactivation recorded from immature and matured hiPSC-CMs at pH 7.4 and acidic pH 6.3. Data were compared using a two-way ANOVA and a two tailed Mann-Whitney test. Errors bars represent SEM. N-value = 3, n value ≥ 9. ****P < 0.0001, ***P = 0.0008, **P = 0.0024, and *P < 0.05.

Next, we normalized the magnitude of IKr at pH 6.3 to the maximum IKr magnitude recorded at pH 7.4 for the same cell (Fig. 2B-E). Remarkably, both steady-state and tail IKr from immature hiPSC-CMs were significantly more sensitive to extracellular acidosis than IKr in matured hiPSC-CMs (Fig. 2B,C, & E). This was particularly true for steady-state IKr, which displayed a roughly two-fold increase in inhibition in immature cells compared to matured cells at 0 through +20 mV (Fig. 2B,C). Similar to our initial recordings (Suppl. Fig. S3B), there is a trend that the time course of IKr deactivation recorded from immature hiPSC-CMs display smaller time constants (117 ± 23 ms at pH 7.4 and 41 ± 11 ms at pH 6.3) than IKr recorded from matured hiPSC-CMs (195 ± 72 ms at pH 7.4 and 80 ± 35 at pH 6.3), Fig. 2G. IKr deactivation at pH 6.3 does not have a slow component of decay.

These results confirm the experimental observations of previous studies on the impact of protons on hERG1 channel activity. And given the distinct deactivation kinetics and proton sensitivities of IKr recorded from immature vs matured hiPSC-CMs, these data also suggest that shifts in hERG1 subunit abundance may mediate the response of native IKr to extracellular acidosis.

### hERG1a and hERG1b expression is dependent upon hiPSC-CM maturation

hERG1 subunit expression is dynamic, varying with development, cell cycle, maturation, and disease states (45,59-63). The slowing of IKr deactivation with maturation suggests an increase in hERG1a relative to hERG1b. The diminished proton sensitivity of IKr in matured cells also suggests that hERG1a is upregulated, as hERG1a was shown to be less sensitive to protons, compared to hERG1b, in CHO cells (42). To examine the expression of the hERG1a and hERG1b subunits in mature and immature hiPSC-CMs, we measured subunit-specific immunofluorescence and mRNA expression levels by qRT-PCR from monolayers cultured on glass or PDMS (Fig. 3).

**Fig. 3.**
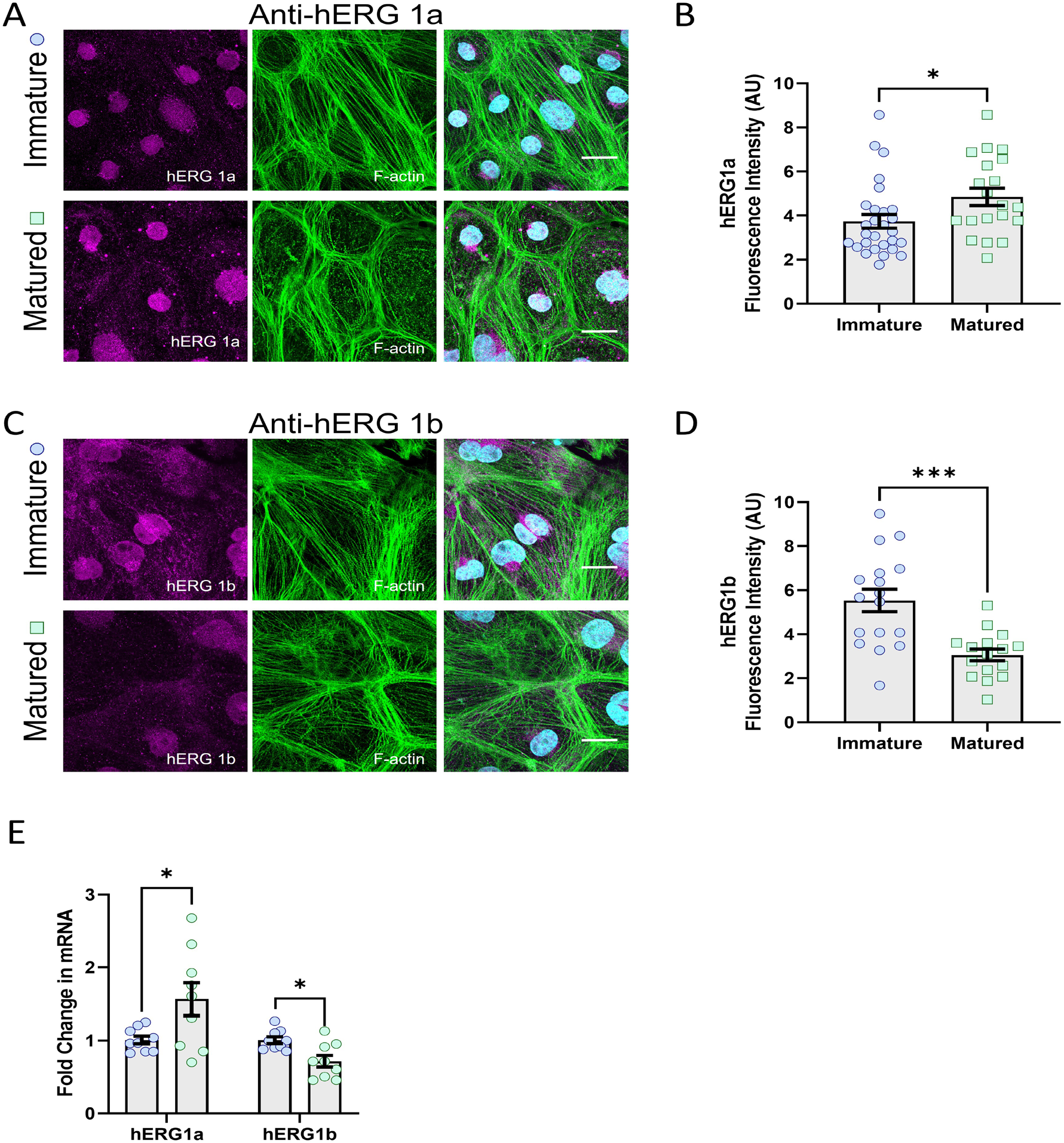
hERG1 subunit abundance in matured and immature hiPSC-CMs. **A)** Representative immunostainings for hERG1a and F-actin. **B)** Quantification of mean hERG1a immunofluorescence from matured and immature hiPSC-CMs. **C)** Representative immunostainings for hERG1b and F-actin. **D)** Quantification of mean hERG1b immunofluorescence from matured and immature hiPSC-CMs. **E)** hERG1a and hERG1b mRNA levels in matured and immature hiPSC-CMs. Data were compared using a two-tailed Mann-Whitney test. Errors bars represent SEM. N-value = 3, n value ≥ 8. ***P = 0.0002, and *P < 0.05.

Consistent with our hypothesis, we found that hERG1a immunofluorescence was significantly increased in matured hiPSC-CM monolayers (4.8 ± 0.4 A.U.) compared to immature hiPSC-CM monolayers (3.7 ± 0.3 A.U.) (Fig. 3A,B). In contrast, hERG1b immunofluorescence was significantly decreased in matured monolayers (3.0 ± 0.3 A.U.) compared to immature monolayers (5.5 ± 0.5 A.U.) (Fig. 4C,D). hERG1a and hERG1b mRNA levels were similarly affected in matured hiPSC CMs compared to immature hiPSC-CMs (1.5 ± 0.2-fold change and 0.7 ± 0.07-fold change in matured cells for hERG1a and hERG1b mRNA levels, respectively), as shown in Figure 3E. These data demonstrate that hiPSC-CM maturation increases hERG1a expression while decreasing hERG1b expression. These data also further support the hypothesis that hERG1 subunit abundance determines IKr proton sensitivity in hiPSC-CMs.

**Fig. 4.**
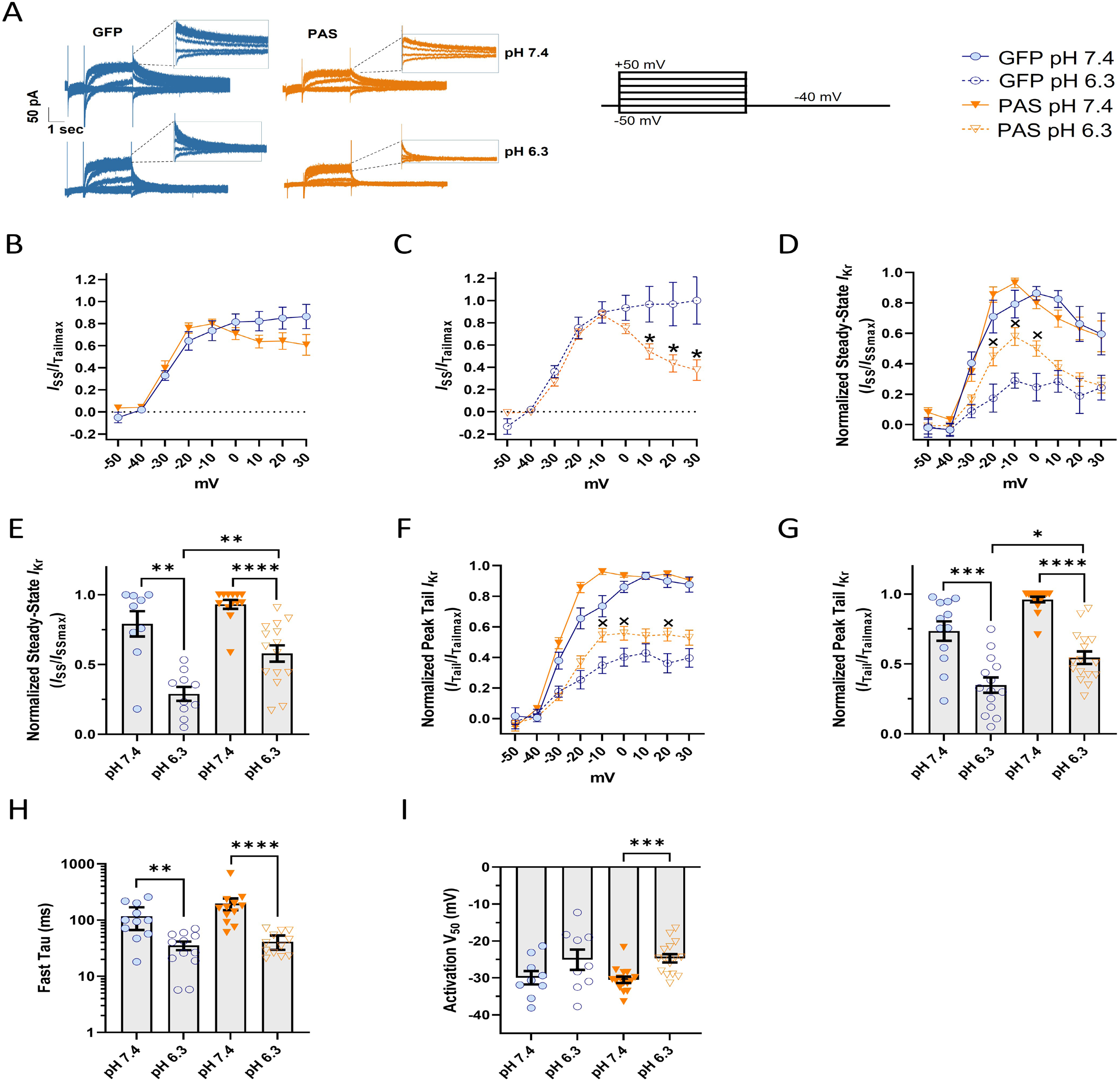
PAS domain expression diminishes IKr proton sensitivity in immature hiPSC-CMs. **A)** Representative IKr traces elicited by the protocol below from PAS (orange) and GFP (blue)-transduced hiPSC-CMs at pH 7.4 and pH 6.3. **B–C)** Steady-state I-V relationships normalized to the maximum peak tail IKr from immature hiPSC-CMs expressing either GFP or PAS, at pH 7.4 and 6.3, respectively. **D)** Steady state IKr at pH 7.4 and 6.3 in immature hiPSC-CMs overexpressing PAS or GFP. **E)** Normalized steady-state current densities at 10 mV. **F)** Peak-tail IKr levels measured at pH 7.4 and 6.3 in immature hiPSC-CMs overexpressing PAS (orange) or GFP (blue). **G)** Normalized tail current densities at 10 mV. **H)** Deactivation kinetics from GFP and PAS-expressing hiPSC-CMs. **I)** Voltage-dependence of activation (V1/2) in immature and matured hiPSC-CMs at pH 7.4 and pH 6.3. The symbol × represent statistical differences between PAS and GFP-transduced hiPSC CMs at pH 6.3. Data were compared using a two-way ANOVA and a two tailed Mann-Whitney test. Errors bars represent SEM. N-value = 3, n value ≥ 10. ****P < 0.0001, ***P = 0.0008, **P < 0.0036, and *P < 0.05.

### PAS expression reduces IKr proton sensitivity in immature hiPSCCMs

Defining the regulatory elements of hERG1 subunits as they pertain to responses to acidosis is a necessary step toward understanding the functional adaptation and impairment of native cardiomyocytes during developmental and pathological processes. Our study revealed that proton inhibition of IKr is enhanced in immature hiPSC-CMs, where hERG1b expression is upregulated. These data suggest that hERG1b expression promotes proton inhibition of IKr. To test this hypothesis, we overexpressed a polypeptide identical to the hERG1a PAS domain in immature hiPSC-CMs (Fig. 4). This technique has been used in heterologous expression systems (24,64) and hiPSC-CMs (16) to mask the impact of hERG1b on heteromeric channel gating. When overexpressed, the PAS polypeptide fills the open receptor site left by the abbreviated hERG1b N-terminal domain (24,64), and thereby transforms heteromeric hERG1a/1b channel gating to a phenotype indistinguishable from homomeric hERG1a channels.

To validate that the PAS polypeptide was appropriately modifying native hERG1 channel function, we first quantified the magnitude of rectification of GFP and PAS-transduced cells. The hERG1a PAS domain promotes inactivation and thereby enhances rectification (20,24), thus IKr recorded from PAS-transduced cells should display increased rectification. We normalized steady-state currents to the maximum peak tail current recorded from the same cell to quantify the magnitude of current inhibition at positive potentials (rectification). As predicted, PAS-transduced cells displayed enhanced rectification of steady-state currents at both pH 7.4 and pH 6.3, compared to GFP-transduced controls (Fig. 4B,C). These data demonstrate that the overexpressed PAS domain is modifying the function of the extant hERG1 channels at both pH 7.4 and pH 6.3. Remarkably, the degree of rectification observed in immature hiPSC-CMs expressing PAS was comparable to that seen in matured hiPSC-CMs (Suppl. Fig. 4).

As expected, pH 6.3 significantly inhibited IKr magnitude (Fig. 4A,D-G), accelerated IKr deactivation (Fig. 4H), and depolarized the voltage-dependence of IKr (Fig. 4I) in both PAS-transduced and GFP-transduced controls. Consistent with our hypothesis, PAS polypeptide overexpression significantly reduced IKr inhibition by protons, compared to GFP controls (Fig. 4A,D-G). At pH 6.3, normalized steady-state IKr was reduced by only 40 ± 6% in PAS-transduced cells compared to 65 ± 9% in GFP-transduced cells, at -10 mV (Fig. 4D & E). Tail IKr was reduced by 44 ± 8% and 54 ± 8% for PAS and GFP-transduced cells, respectively (Fig. 4F & G). In fact, the magnitude of proton inhibition of IKr in PAS-transduced cells was comparable to that observed in our matured hiPSC-CMs (cf. Fig. 2). Together, our findings shed light on how the hERG1a PAS domain, in addition to modulating the kinetic properties of channel gating, plays an important role in the response of hERG1 channels to extracellular acidosis. Finally, these data also demonstrate that the relative abundance of hERG1a and hERG1b subunits influences the magnitude of IKr inhibition by extracellular protons.

## DISCUSSION

The present study investigates the impact of extracellular acidosis on native IKr recorded from hiPSC CMs. First, as demonstrated by others (56), the data presented herein validate that hiPSC-CMs cultured on a soft Matrigel-coated PDMS substrate displayed electrophysiological features consistent with enhanced cardiac maturation, including hyperpolarized resting membrane potentials and increased action potential amplitude, when compared to cells plated on Matrigel-coated glass. Additionally, IKr recorded from PDMS-matured hiPSC-CMs was less sensitive to extracellular protons compared IKr recorded from immature hiPSC-CMs. Finally, the decrease in proton sensitivity between immature and matured cells was mediated by an increase in the relative abundance of hERG1a and hERG1b subunits at the cell surface membrane.

### Proton Modulation of hERG1

The impact of external protons on hERG1a has been well-described in heterologous expression systems, with distinct effects on single channel conductance and gating (31,33,34,37,41). Surprisingly, the impact of protons on native IKr is poorly described. This is of particular importance because native cardiac hERG1 channels comprise both hERG1a and hERG1b subunits (15,16,18), and are modulated by other potential accessory subunits (e.g. KCNE1 and KCNE2) and interacting proteins (e.g. KvLQT1) (65-68).

Here we demonstrated that reduced extracellular pH inhibited current density and depolarized the voltage dependence of IKr recorded from hiPSC-CMs. These data are consistent with other work on native IKr (27,69). Interestingly, our study demonstrated that the degree of IKr inhibition by protons correlated with the relative abundance of the hERG1b subunit. Work in CHO cells has also demonstrated that inhibition of hERG1 conductance by protons is more pronounced in channels that contain the hERG1b subunit, compared hERG1a homomeric channels (42). In this study, proton inhibition of IKr was greatest in immature iPSC-cardiomyocytes, where hERG1b was upregulated. The enhanced inhibition by protons was then abolished by increasing the number of PAS domains per channel, effectively transforming hERG1b subunits into hERG1a subunits. Importantly, the time course of deactivation is an additional marker of PAS activity, where PAS-deficient heteromeric hERG1a/1b channels display faster deactivation compared to homomeric hERG1a channels. However, because of the dramatic accelerating effects of protons on deactivation, it is not a reliable marker of PAS action at reduced pH. Accordingly, our data demonstrate that hERG1 subunit stoichiometry mediates proton inhibition of IKr in hiPSC-CMs.

It is unclear how hERG1b selectively enhances proton inhibition of channel conductance without altering the impact of protons on channel gating. This is somewhat surprising given the pronounced accelerating effects that hERG1b has on hERG1 channel gating, particularly deactivation (14,15,20). hERG1 has proton binding sites at the pore and voltage-sensing domains that modulate conductance and gating, respectively, with different pH sensitivities (31,33,34,37,41,70). At the voltage sensing domain, mutating a trio of aspartates to alanines (D456A/D460A/D509A) disrupts proton modulation of channel gating (33,70). Proton block, however, is critically dependent upon residues E575 and H578 at the hERG1 pore turret, where the combined mutations E575Q and H578N abolish proton block without affecting proton modulation of deactivation (41). Although they are located on the outer circumference of the hERG1 pore, these proton binding sites (at least E575 and H578) alter the electrostatic environments in and around the selectivity filter (41). It was proposed that the outer hERG pore near the selectivity filter is somewhat flexible (71-73), underlying inactivation and possibly providing a mechanism to transmit protonation of E575 and H578 to changes in hERG1 channel conductance and open time (41,72,74). Nonetheless, this must be approached with caution because only the E575 side chain was shown in the hERG1 cryo-EM structure to directly interact with residues that connect to the selectivity filter (72). Because the residues involved in proton sensitivity are found in both hERG1a and hERG1b, it is possible that based on the cryo-EM structure, the greater effect we observed on cells preferentially expressing hERG1b was due to indirect/allosteric consequences of the unique hERG1b N-terminus that favor exposure of E575 and H578 to protons. Intracellular acidosis does not affect hERG1a homomeric channels (38), but we cannot rule out that the short hERG1b N-terminus may expose intracellular proton binding sites, otherwise occluded by the hERG1a PAS domain.

### Protons in IKr-Mediated Cardiac Dysfunction

Cardiac acidosis occurs under a number of physiological and pathophysiological conditions. Two conditions, ischemic heart disease and sudden infant death syndrome (SIDS), are particularly affiliated with hERG1 dysfunction. In chronic cardiac dysfunction, i.e. heart failure, native IKr is significantly downregulated (75,76) and the relative abundance of hERG1b to hERG1a is increased (51). These changes in IKr occur alongside the downregulation of other major K+ currents: IKs, Ito, and IK1 (77-79). The reduced IK contributes collectively to the reduced repolarization reserve, prolonged action potential duration, and overall heightened arrhythmogenic potential in the failing myocardium. Our data suggest that a relative increase in hERG1b in the failing heart would also enhance IKr sensitivity to protons during ischemic events. And although hERG1b homomeric channels may not exist in adult hearts – hERG1b subunits preferentially associate with hERG1a – (80,81), the relative expression of hERG1a and hERG1b subunits appears heterogeneous in cardiac tissue (17,48,82). In this regard, regional variation in hERG1 isoform abundances could facilitate heterogeneity of repolarization and arrhythmogenesis during acidosis.

### hERG1 subunits in neonatal and fetal demise

KCNH2 variants have long been linked with sudden infant death syndrome (SIDS) (3,7,13). Respiratory acidosis is one hypothesis proposed to explain the association of stomach sleeping with SIDS (8,83). Interestingly, hERG1a mRNA is upregulated and hERG1b mRNA is downregulated in adult human cardiac tissue compared to fetal cardiac tissue (3). These molecular data combined with our electrophysiological data suggest that upregulated hERG1b in the immature heart could promote proton inhibition of IKr during respiratory acidosis, and thereby contribute to SIDS. The shifts in subunit abundance during maturation also predict that the pathophysiological impact of hERG1b-specific mutations would be greatest in the immature heart and vice versa for hERG1a-specific mutations. Indeed, the only two hERG1b-specific mutations identified to date were a case of intrauterine fetal death, R25W, (3) and an 8-year old girl, A8V, (20). Interestingly, similarly to other mutations found in SIDS cases (R273Q and R954C/K897T), the mutation R25W generates a profound reduction in current density when expressed as heterotetramers with the hERG1a subunit (3,4,9).

For normal heart function, these two hERG1 subunits must be functionally expressed. Changes in the abundance of hERG1a or hERG1b can cause proarrhythmic events (16,20,46). Clearly, there is a link between LQTS2 and intrauterine fetal death (3,84-86). hERG1 channel variants that have been solely linked with SIDS have the potential to be LQTS variants. And it is possible that LQTS cases are being disguised under the SIDS umbrella, settinga precedent forfuture research into the role of cardiac channelopathies.

### Subunit-Selective Modulators

Protons are not the only factor shown to differentially modulate homomeric and heteromeric hERG1 channels. Several studies have demonstrated that subunit abundance, and its impact on gating, mediates the channel’s response to a subset of clinically relevant drugs (20,87,88). Additionally, ANP and cGMP perfusion were both shown to selectively inhibit hERG1b-containing channels heterologously expressed in HEK293 cells (89). In the same study, the authors demonstrated that cGMP inhibited IKr recorded from atrial but not ventricular murine myocytes, suggesting that mERG1b (mouse ERG1b) was more expressed in atrial than in ventricular murine tissue (89). Assuming that mERG1b is primarily expressed in the atria, and based on findings from computational modelling indicating that the gain-of-function mutations L532P and N588K cause a higher and earlier peak of IKr during atrial APs and lead to rotor formation (90), we postulate that ERG1b subunit expression may play a key role in atrial fibrillation (91). In contrast, hERG1b shows a protective effect against oxidative inhibition, presumably by regulating access to a key residue in the channel’s C-linker domain C723 (hERG1a numbering). Roughly two thirds of the protective effect from hERG1b was attributable to the subunit’s acceleration of channel deactivation (92).

The fact that hERG1b is upregulated in “immature” hiPSC-cardiomyocytes underscores the need for increased understanding of mechanisms regulating hERG1 subunit abundance. Drugs that preferentially target hERG1 isoforms may be one approach to overcome obstacles in treating disorders in the heart and other tissues where hERG1 is a contributing factor. The two hERG1 isoforms are expressed in distinct ratios and contribute differently to the maintenance of hERG1 currents in tissues where hERG1 is functional. For example, while hERG1b is expressed at lower levels in the human heart, it is the predominant isoform in tumor cells (93). In B cells and T-cell lineage, while hERG1b is upregulated, the other isoform, hERG1a, is downregulated (94). Therefore, identifying the mechanisms that control hERG1 subunit abundance could improve clinical therapies in diseases throughout the body.

## Conclusion

The experimental data presented herein show for the first time the effects of extracellular acidosis in hiPSC-cardiomyocytes that express one subunit preferentially, either hERG1a in PDMS-plated cells or hERG1b in glass-plated cells. And although the exact tetrameric conformation of native hERG1 channels remains elusive, these findings provide insight into the response of adult and immature cardiomyocytes to an acidic environment.

### Limitations

Here, we report data from experiments conducted in immature and matured hiPSC CMs. While our studies demonstrate the impact of extracellular acidosis in a human cardiomyocyte model, hiPSC-CMs still cannot recapitulate the chamber-specific or layer-specific electrical phenotypes of intact cardiac tissue. And though tools to enhance hiPSC-CM maturation have improved, matured hiPSC-CMs still display significant differences compared with the adult ventricular cardiomyocyte. Thus, the magnitude of the effects observed in this manuscript should be extrapolated to the adult myocardium with caution. Additionally, native IKr magnitudes are relatively small, particularly at pH 6.3, which increases experimental variability. Nonetheless, these data provide important insight into the triggers of IKr dysfunction during extracellular acidosis.

## MATERIALS AND METHODS

### Stem Cell Culture and Cardiac Differentiation

Human iPS cells were cultured and differentiated into cardiomyocytes using the GiWi protocol, as described (52). Briefly, stem cells were seeded on Matrigel-coated plasticware with iPS-brew medium. Spontaneous differentiation was removed, and cells were passed at 70% confluence. At the day of cell passage, cells were re-seeded to continue the line, or to grow monolayers for cardiac-directed differentiation. 4 × 105 cells were plated into each well of a 6-well plate and cultured to ∼80% confluence for treatment with GSK3 inhibitor and induction of mesodermal differentiation (day 0). Following mesodermal differentiation, cells were treated with a Wnt inhibitor for induction of cardiac mesoderm (day 2). On day 4, Wnt inhibitor was removed to direct the cells into cardiac progenitor cells. iPSC-cardiomyocytes with autonomous contractility emerged eight to ten days after initiation of cardiac-directed differentiation. The iPSC-cardiomyocytes were cultured until 20 days after initiation of differentiation and purified using by magnetic-beads assisted isolation with an iPSC-Derived Cardiomyocyte Isolation Kit, human (Miltenyi Biotec, USA). Purified iPSC-cardiomyocytes were then plated on either Matrigel-coated glass coverslips (immature cells) or Matrigel-coated polydimethylsiloxane (matured cells) for seven days before completing experiments.

### Stem Cell Culture and Cardiac Differentiation

We cultured and differentiated human iPS and ES cells into cardiomyocytes using the GiWi protocol, as described (42). Briefly, we seeded stem cells on Matrigel-coated plasticware with iPS-brew medium. We checked media daily to remove spontaneous differentiation and passed the cells at 70% confluence. At the day of cell passage, we re-seeded cells to continue the line, or seeded the cells to grow monolayers for cardiac-directed differentiation. We plated 4 × 105 cells into each well of a 6-well plate and cultured them to ∼80% confluence for treatment with GSK3 inhibitor for induction of mesodermal differentiation (day 0, D0). Following mesodermal differentiation, we treated cells with a Wnt inhibitor for induction of cardiac mesoderm (D2). On D4 we removed Wnt inhibition to direct the cells into cardiac progenitor cells. Cardiomyocytes with autonomous contractility emerged eight to ten days after initiation of cardiac-directed differentiation. We cultured the cardiomyocytes until 20 days after initiation of differentiation, and isolated purified cardiomyocytes by magnetic-beads assisted isolation with an iPSC-Derived Cardiomyocyte Isolation Kit, human (Miltenyi Biotec, USA) following the manufacturer’s recommendations. We plated the purified cardiomyocytes on Matrigel-coated coverslips for seven days before completing experiments.

### RT-qPCR

For quantitative evaluation of the steady-state mRNA expression in hiPSC-CM cultures, total RNA was prepared using the RNeasy Mini Kit (Qiagen), including DNAse treatment. 300 ng of RNA were reverse transcribed and converted to cDNA with oligo(dT)12–18 primers using reverse transcriptase according to the manufacturer’s specifications, M-MLV Reverse transcriptase (Cat # 28025-013, Invitrogen). Quantitative PCR was performed using IDT Mastermix (Cat # 1055772, ThermoFisher) and TaqMan assay primers (Cat # 4331182 and 43513752, 10 μM; ThermoFisher) for KCNH2, 1a and 1b isoforms. The PCR condition consisted of 95°C for 30 secs, followed by 39 cycles of 95°C for 3 secs and 60°C for 20 secs, followed by melting-curve analysis to verify the correctness of the amplicon.

The samples were analyzed in technical triplicates using the primers included in the TaqMan Assay system (Invitrogen) and run in a Biorad C1000 Touch Thermal Cycle CFX96 (Applied Biosystems). The expression of the mRNA of the gene of interest relative to the internal control GAPDH in samples from immature and matured hiPSC CMs was calculated by the ΔΔCT method, based on the threshold cycle (CT), as fold change = 2^−(ΔΔCT), where ΔCT = CTgene of interest − CTGAPDH and ΔΔCT = ΔCTMatured hiPSC-CMs – ΔCTImmatured hiPSC-CMs. From each experiment, the cDNA of 3 cell culture wells were measured as biological replicates of each cell maturation state. Each cell culture well was measured from at least 3 separate cardiomyocyte differentiation.

### Immunocytochemistry

hiPSC-CMs were seeded either on glass or PDMS and fixed with 4% paraformaldehyde/PBS for 15 min. Then, hiPSC-CMs were washed 5 min with PBS and blocked with block solution (PBS + 1% BSA + 0.5% Triton X + 10% Goat Serum secondary antibodies) for 1 h. Incubation with primary antibodies was done in block solution for overnight at 4°C. The next day, to washout the excess of primary antibody, hiPSC-CMs were washed 3×5min with PBS. Next, secondary antibodies in block solution (without Triton X) were added to each slip and incubated for 1 h in the dark at room temperature. hiPSC-CMs were kept in dark, washed with PBS 3 × 5 min, and mounted with ProLong Gold antifade reagent (Thermo Fisher) and a coverslip. Both primary and secondary antibodies were diluted in block solution (without Triton-X).

Differentiated cardiac lines were validated using immunocytochemistry targeting actin (phalloidin, cat #A12379 ThermoFisher) to display the cardiac sarcomeric organization, and patch clamp electrophysiology measuring cardiac IKr, indicative of hERG1 expression. Phalloidin-488 comes with a fluorophore conjugated so no secondary Ab incubation was needed. To target the hERG1a isoform, iPSC-cardiomyocytes were immunolabeled with a 1:200 dilution of the primary antibody #ALX-215-050-R100 (Enzo Life Sciences). To target the hERG1b isoform, the primary antibody #ALX-215-051-R100 was used in a 1:200 dilution (Enzo Life Sciences). In both cases, a 1:250 dilution of secondary antibody goat anti-rabbit Alexa Fluor 647 (#4050-31, Southern Biotec) was used. The nuclei were labeled using 1:1000 dilution of DAPI (1μg/ml) for 15 minutes (ThermoScientific, Cat. #62248). Immunostained preparations were analyzed by confocal microscopy, using a confocal microscope (Zeiss 880) to determine protein localization.

### Electrophysiology

Standard patch-clamp techniques were used to measure both action potential clamp waveform and IKr. All recordings were completed at physiological temperature (37±1°C) using whole-cell patch clamp with an IPA® Integrated Patch Amplifier run by SutterPatch® (Sutter Instrument) and Igor Pro 8 (Wavemetrics). Leak subtraction was performed off-line based on measured current observed at potentials negative to IKr activation. The inter-pulse duration for all recordings was 10 seconds where cells were at -40 mV.

Data were sampled at 5 kHz and low-pass filtered at 1 kHz. Cells were perfused with extracellular solution containing (in mM): 150 NaCl, 5.4 KCl, 1.8 CaCl2, 1 MgCl2, 15 glucose, 10 HEPES, 1 Na-pyruvate, and titrated to pH 7.4 using NaOH. Recording pipettes had resistances of 2–5 MΩ when backfilled with intracellular solution containing (in mM): 5 NaCl, 150 KCl, 2 CaCl2, 5 EGTA, 10 HEPES, 5 MgATP and titrated to pH 7.2 using KOH. Intracellular solution aliquots were kept frozen until the day of recording. We kept the intracellular solution on ice during recordings and discarded it 2–3 hours post-thaw.

To isolate IKr, all protocols were completed before and after extracellular perfusion of 2μM of the IKr-specific blocker, E-4031. To inactivate sodium currents, a 100-ms step to 40 mV was applied before the any IKr recordings. To assess the voltage dependence of IKr activation, cells were stepped from a holding potential of -40 mV to a three second pre-pulse between -50 and +50 mV in 10 mV increments. Tail currents were then measured during a -40 mV, 3 second test pulse. Peak tail current was normalized to cellular capacitance, plotted as a function of pre-pulse potential, and fitted with the following Boltzmann equation:

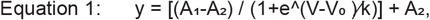

where A1 and A2 represent the maximum and minimums of the fit, respectively, V is the membrane potential, V0 is the midpoint, and k is the slope factor. The time course of IKr deactivation was assessed by fitting current decay during the test pulse with a double exponential function:

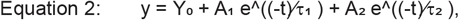

where Y_0_ is the asymptote, A_1_ and A_2_ are the relative components of the fast and slow time constants τ_1_ and τ_2_, respectively. The magnitude of IKr rectification was quantified by dividing the average IKr during the final 10 ms of each step pulse by the maximum peak outward tail current evoked at -40 mV. Repolarizing charge was calculated by integrating IKr recorded during a voltage protocol that mimics a human ventricular action potential (53).

## Statistical Analysis

Analysis was completed using Prism 8 (GraphPad) and Igor Pro 8 (Wavemetrics). Values were first tested for normality (Shapiro-Wilk test) before statistical evaluation. All data are reported as mean ± SEM and were compared using a non-parametric Mann-Whitney test or 2-way ANOVA with a Bonferroni post-hoc test, where applicable. Statistical significance was taken at p < 0.05. Data points greater than two times the standard deviation were termed outliers and excluded from analysis. Unless stated otherwise, the number n of observations indicated reflects the number of hiPSC-CMs recorded from each cell line from at least 3 differentiations. All experiments were performed as a single-blind study to avoid sources of bias.

## ACKNOWLEDGMENTS

This research was supported by NIH/NHLBI R00HL133482 to DKJ; Training Program in Translational Cardiovascular Research and Entrepreneurship 5 T32 HL 125242-7, and Pharmacological Sciences Training Program T32-GM007767 to CUU.

