## Supplemental figures for "hERG1 channel subunit composition mediates proton inhibition of IKr in hiPSC-CMs" (from another version of this preprint: version 1, 2022-08-27)

### SUPPLEMENTARY FIGURES

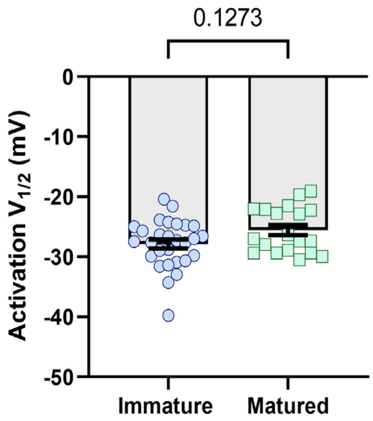


**Suppl. Fig 1. Voltage-dependence of activation V1/2.** There was no significant difference in the V1/2 for IKr activation recorded from immature hiPSC-CMs versus matured hiPSC-CMs. Two tailed Mann-Whitney test. Errors bars represent SEM. N value = 3, n value >10.

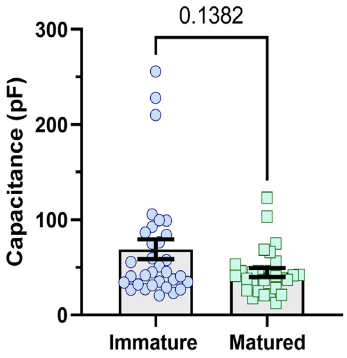


**Suppl. Fig 2. hiPSC-CM Capacitance.** There was no statistically significant difference in the cellular capacitance of immature and matured hiPSC-CMs. Two tailed Mann-Whitney test. Errors bars represent SEM. N-value = 3, n value >10.

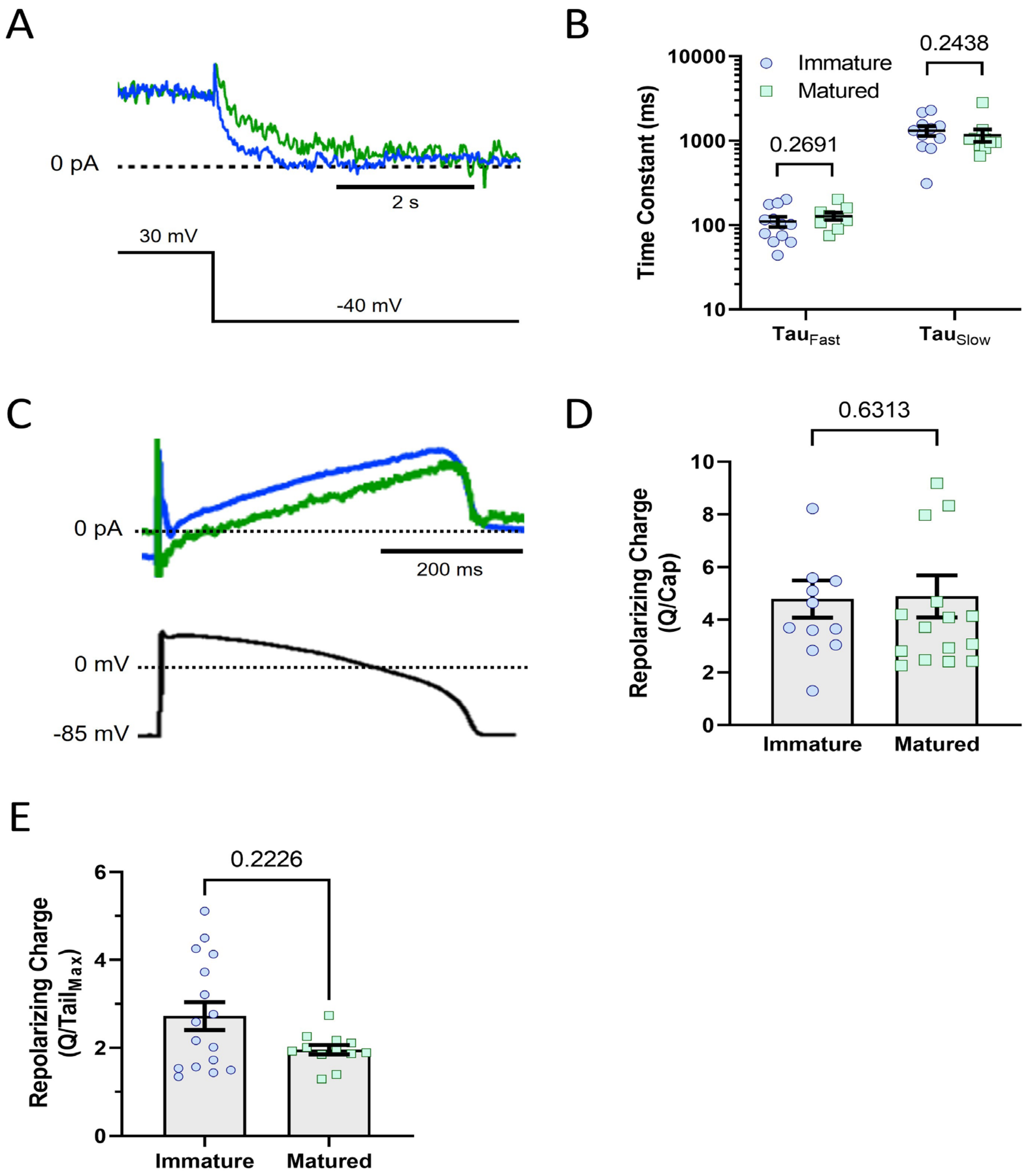


**Suppl. Figure 3. Effect of cell maturation on IKr deactivation.** A) Tail IKr traces recorded from immature (blue) and matured (green) hiPSC-CMs at -40 mV. B) Corresponding deactivation time constants for IKr recorded from hiPSC-CMs cultured on glass (immature) versus PDMS (matured). C) Representative IKr traces recorded from immature (blue) and matured (green) hiPSC-CMs using a ventricular AP clamp protocol. D) Repolarizing charge normalized to the cell capacitance in matured and immature hiPSC-CMs. E) Repolarizing charge normalized to the peak-tail IKr of the same cell recorded from matured and immature hiPSC-CMs. Two tailed Mann-Whitney test. Errors bars represent SEM. N-value = 3, n value >10.

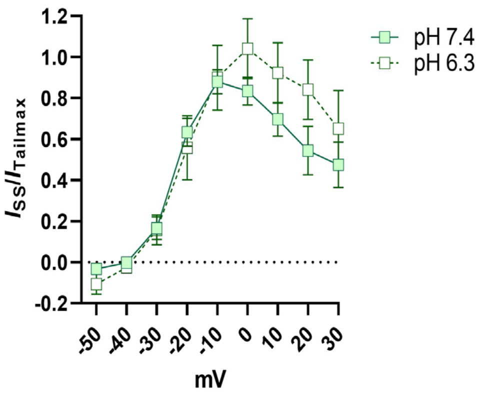


**Suppl. Figure 4. Degree of IKr rectification in matured hiPSC-CMs.** Steady-state I-V relationships normalized to the maximum peak tail IKr recorded from the same cell for matured hiPSC-CMs at pH 7.4 (solid symbols) and 6.3 (open symbols), respectively. Errors bars represent SEM. N-value = 3, n value >9.
